# Control of three-carbon amino acid homeostasis by promiscuous importers and exporters in *Bacillus subtilis*: Role of the sleeping beauty family of amino acid exporters

**DOI:** 10.1101/2023.12.18.572190

**Authors:** Robert Warneke, Christina Herzberg, Richard Daniel, Björn Hormes, Jörg Stülke

## Abstract

The Gram-positive model bacterium *Bacillus subtilis* can acquire amino acids by import, *de novo* biosynthesis, or by degradation of proteins and peptides. The accumulation of several amino acids inhibits growth of *B. subtilis*, probably due to misincorporation into cellular macromolecules such as proteins or peptidoglycan or due to interference with other amino acid biosynthetic pathways. Here, we studied the adaptation of *B. subtilis* to toxic concentrations of the three-carbon amino acids L-alanine, β-alanine, and 2,3-diaminopropionic acid as well as glycine. Resistance to the non-proteinogenic amino acid β-alanine, which is a precursor for the vitamin B5 and thus for coenzyme A biosynthesis is achieved by mutations that either activate a cryptic amino acid exporter, AexA (previously YdeD), or inactivate the amino acid importers AimA, AimB (previously YbxG), and BcaP. The *aexA* gene is very poorly expressed under most conditions studied. However, mutations afecting the transcription factor AerA (previously YdeC), can result in strong constitutive *aexA* expression. AexA is the founding member of a conserved family of amino acid exporters in *B. subtilis*, which are all very poorly expressed. Therefore, we suggest to call this family “sleeping beauty family of amino acid exporters”. 2,3-Diaminopropionic acid can also be exported by AexA, and this amino acid also seems to be a natural substrate of AerA/ AexA, as it can cause a slight but significant induction of *aexA* expression, and AexA also provides some natural resistance towards 2,3-diaminopropionic acid. Moreover, our work shows how low specificity amino acid transporters contribute to amino acid homeostasis in *B. subtilis*.

**IMPORTANCE:** Even though *B. subtilis* is of of the most-studied bacteria, amino acid homeostasis in this organism is not fully understood. We have identified import and export systems for the C2 and C3 amino acids. Our work demonstrates that the responsible amino acid permeases contribute in a rather promiscuitive way to amino acid uptake. In addition, we have discovered AexA, the first member of a family of very poorly expressed amino acid exporters, that we call “sleeping beauty amino acid exporters”. The expression of these transporters is typically triggered by mutations in corresponding regulator genes that are acquired upon exposure to toxic amino acids. These exporters are ubiquitous in all domains of life. It is tempting to speculate that many of them are not expressed until the cells experience a selective pressure by toxic compounds and that the protect the cells from rare but potentially dangerous accounters with such compounds.

## INTRODUCTION

Amino acids are major building blocks of all living cells. In addition to the proteinogenic amino acids there are also non-proteinogenic amino acids that may have important functions for the cell. In bacteria, meso-diaminopimelic acid and ornithine as well as D-glutamate and D-alanine are essential building blocks of the peptidoglycan cell wall (1). β-alanine, the only naturally occuring β-amino acid, is required as a precursor for the biosynthesys of coenzyme A (2, 3). Citrulline, ornithine, and γ-aminobutyrate can be used as carbon and nitrogen sources, and the latter two serve also as inducers for transcription factors in *Bacillus subtilis* (4, 5, 6). Finally, citrulline serves as a compatible solute to protect the cell from osmotic stress (7).

While amino acids are not only important to the cells, they can also be harmful. Often, the addition of amino acids results in growth inhibition. This is the case for some of the natural proteinogenic and non-proteinogenic amino acids but also for artificial analogs (8). As an example, serine can give rise to α-aminoacrylate and to the highly reactive, and therefore toxic, β-hydroxypyruvate through deamination and transamination, respectively (9, 10). The growth inhibition can also be caused by the reactivity of the amino acids leading to the formation of toxic metabolites. In the case of natural amino acids, their toxicity may also be the result of chemical similarity to other amino acids that may result in binding to enzymes of non-cognate biosynthetic pathways and thus inhibition of biosynthesis and subsequent limitation of a different amino acid. This effect is well studied in the case of serine which in *Escherichia coli* can bind to and inactivate the bifunctional enzyme aspartate kinase/homoserine dehydrogenase (ThrA) (11), thus interfering with threonine biosynthesis. The effect of serine interference with threonine synthesis also occurs in *B. subtilis* (12). Non-natural amino acids can also be toxic for the same reasons but in addition they can also be misincorporated into proteins due to the poor discrimination of amino acyl-tRNA synthetases against these amino acid analogs which was never required during evolution of these systems (13).

Bacteria have several mechanisms to protect themselves against the action of growth inhibiting amino acids. For some of them, the bacteria have degradative pathways that can convert the toxic amino acids to non-inhibiting metabolites. However, these degradation mechanisms are often not sufficiently active in wild-type strains and only their mutational activation allows the effective disposal off some of the amino acids, as shown for serine in *B. subtilis* and for *B. subtilis* strains that are sensitive to glutamate (12, 14, 15). In some cases, bacteria have specific protective mechanisms against the misincorporation of D-amino acids into proteins, e.g. by producing a D-aminoacyl-tRNA deacylase that can recycle misaminoacylated D-Tyr-tRNA(Tyr) and some other D-aminoacyl-tRNAs (13). In addition, a few bacteria, including *B. subtilis*, possess even a second tyrosine-specific amino acyl-tRNA synthetase that strongly discriminates against D-tyrosine. However, expression of the corresponding gene requires a mutation that inactivates the transcription repressor (16). Interestingly, the D-aminoacyl-tRNA deacylase is inactive in many *B. subtilis* strains, they thus have to rely on the more selective Tyr-tRNA synthetase (17). A third way to deal with toxic amino acids is their export: This has been described for the leucine analog 4-azaleucine and histidine. In both cases, export by the AzlCD amino acid exporter only becomes possible when the transcription repressor AzlB that normally prevents the expression of the exporter is inactivated (18, 19).

In addition to the active strategies to deal with harmful amino acids, the bacteria can also deny them access to the cell by inactivation of the corresponding transporters. The use of toxic amino acid analogs or the identification of conditions under which a given amino acid becomes growth-inhibiting is a powerful tool for the identification of amino acid transporters. This approach has been used to identify transporters for serine, threonine, proline, glutamate, and asparagine as well as for the toxic inhibitors of amino acid biosyntheses, glyphosate and glufosinate, in *B. subtilis* (12, 15, 20, 21, 22, 23). This strategy, however, has limited application as some amino acids are imported by multiple transporters (19, 24), and thus the simultaneous inactivation of all of them would be required to prevent the uptake. Moreover, in natural ecosystems the loss of the ability to exploit important resources such as amino acids from the environment would certainly compromise the competetiveness of such bacteria.

We have long-standing interest in the control of amino acid homeostasis in the model bacterium *B. subtilis* (5, 19, 25, 26, 27). Although *B. subtilis* is one of the best studied bacteria, the transporters for several amino acids as well as the substrates of several potential amino acid transporters have not yet been elucidated (24, 28, 29). With the exception of γ-aminobutyrate (30), the transport of non-proteogenic amino acids has not been studied in *B. subtilis*. In this work, we have studied the homeostasis of C3 amino acids in *B. subtilis*, α-L-alanine, β-alanine, and 2,3-diaminopropionic acid. Using the fact that all three amino acids inhibit growth of *B. subtilis*, we identified the underlying importers and exporters. All three amino acids are transported into the cell by the broad-range amino acid transporter AimA. In addition, α-L-alanine and β-alanine are taken up by AlaP, whereas, diaminopropionic acid can also be transported by BcaP and YbxG. The latter protein also acts as a broad-range amino acid permease, which we therefore rename AimB (<u>a</u>mino acid <u>im</u>porter B). Again, all three C3 amino acids can be exported by the poorly expressed amino acid exporter YdeD, which we rename AexA (<u>a</u>mino acid <u>ex</u>porter A). AexA is the founding member of a family of “sleeping beauty” amino acid exporters. Expression of the corresponding *aexA* gene can either be induced by the presence of 2,3-diaminopropionic acid or by mutational activation of the transcription factor YdeC, which we rename AerA (<u>a</u>mino acid <u>e</u>xport regulator A).

## RESULTS

### Three-carbon amino acids inhibit the growth *B. subtilis*

To test how *B. subtilis* can cope with aminated propionic acid derivatives, we tested growth of the wild type strain 168 in the presence of L-α-alanine, β-alanine, and 2,3-diaminopropionic acid (Fig. 1A) in LB complex medium and C minimal medium. As shown in Fig. 1B, the strain grew on both media in the absence of any added amino acid. Similarly, the bacteria were efficiently plated on LB medium irrespective of the presence of the amino acids. In contrast, we observed a concentration-dependent growth inhibition by all three C3 amino acids in minimal medium (Fig. 1C). This inhibition was most severe with diaminopropionic acid were 2 mM caused a nearly total growth inhibition. In contrast, the inhibitive effect was less pronounced for β-alanine and even less for the proteinogenic amino acid L-α-alanine. Since β-alanine is of biological relevance as precursor for the biosynthesis of coenzyme A, we focussed first on this amino acid.

**Figure 1:**
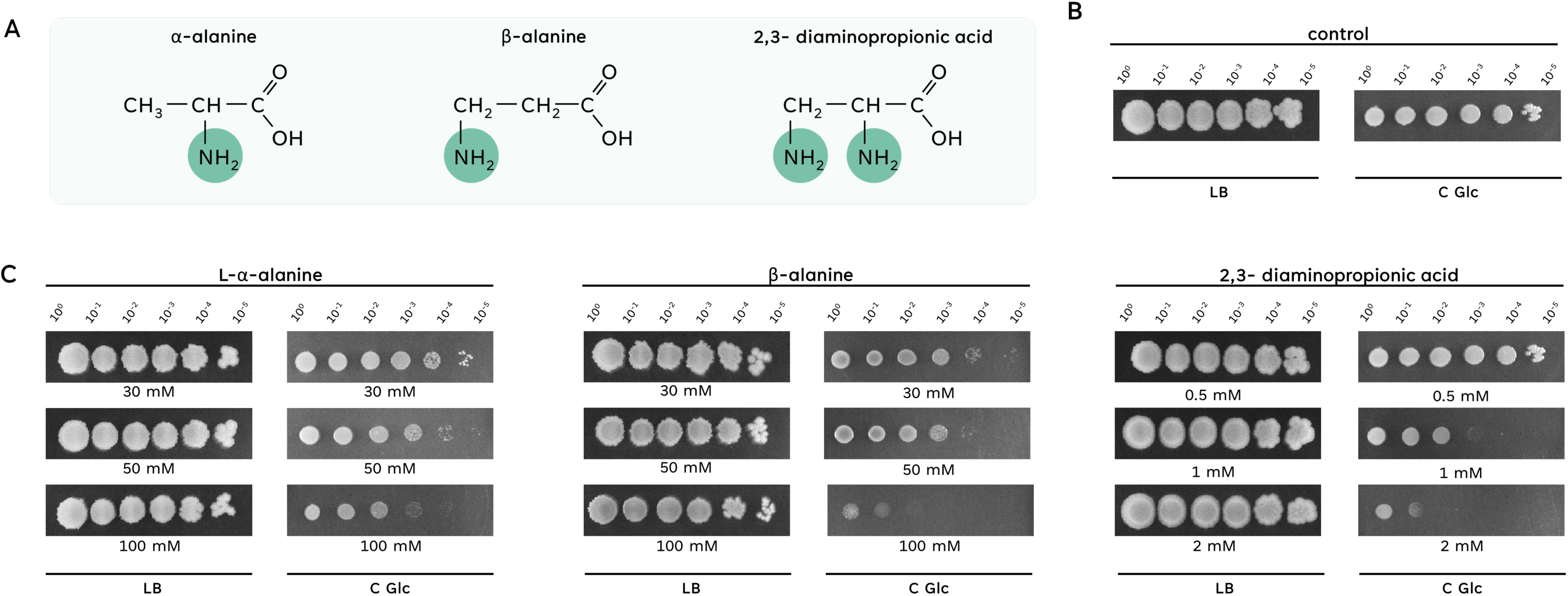
Growth of *Bacillus subtilis* in presence of alanine derivates. **A.** The three alanine derivates a-alanine, β-alanine and 2,3-diaminopropionic acid are displayed. The position of the amino group is highlighted. **B+C.** The growth of the wild type strain 168 was compared on complex medium (LB) and minimal medium (C Glc) in absence or presence of the alanine derivates L-α-alanine, β-alanine and 2,3-diaminopropionic acid. The cells were grown in C-Glc minimal medium or LB complex medium to an OD_600_ of 1.0 and serial dilutions (10-fold) were prepared. These samples were plated on C-Glc minimal plates or LB plates containing 30 mM, 50 mM or 100 mM L-α-alanine or β-alanine, or 0.5 mM – 2 mM 2,3-diaminopropionic acid. The plates were incubated at 37°C for 48 h.

### A suppressor screen identifies a regulator involved in tolerance to β-alanine

To get more insights into the toxicity of β-alanine, we isolated suppressor mutants that were able to grow at increased β-alanine concentrations. In a first step, mutants were isolated at 30 mM β-alanine. Of six isolated suppressor mutants that were viable in the presence of β-alanine, two were subjected to whole genome sequencing. In both mutant strains, we identified single point mutations in the *aerA* (*ydeC*) gene, encoding a so far unkown transcription factor of the AraC family (31). In strain GP4117, Tyr-179 is replaced by an aspartate residue, and in GP4119, Leu-122 is replaced by a tryptophan. The fact that *aerA* was the only affected gene in both cases, suggested an important role for this gene in the acquisition of resistance to β-alanine. We therefore sequenced the *aerA* allele in the remaining four mutants and found that all had the Y179D substitution as in GP4117. Thus, the potential AraC-type transcription factor plays a key role in the adaptation of *B. subtilis* to β-alanine (see Fig. 2 for an overview on all isolated suppressor mutants).

**Figure 2:**
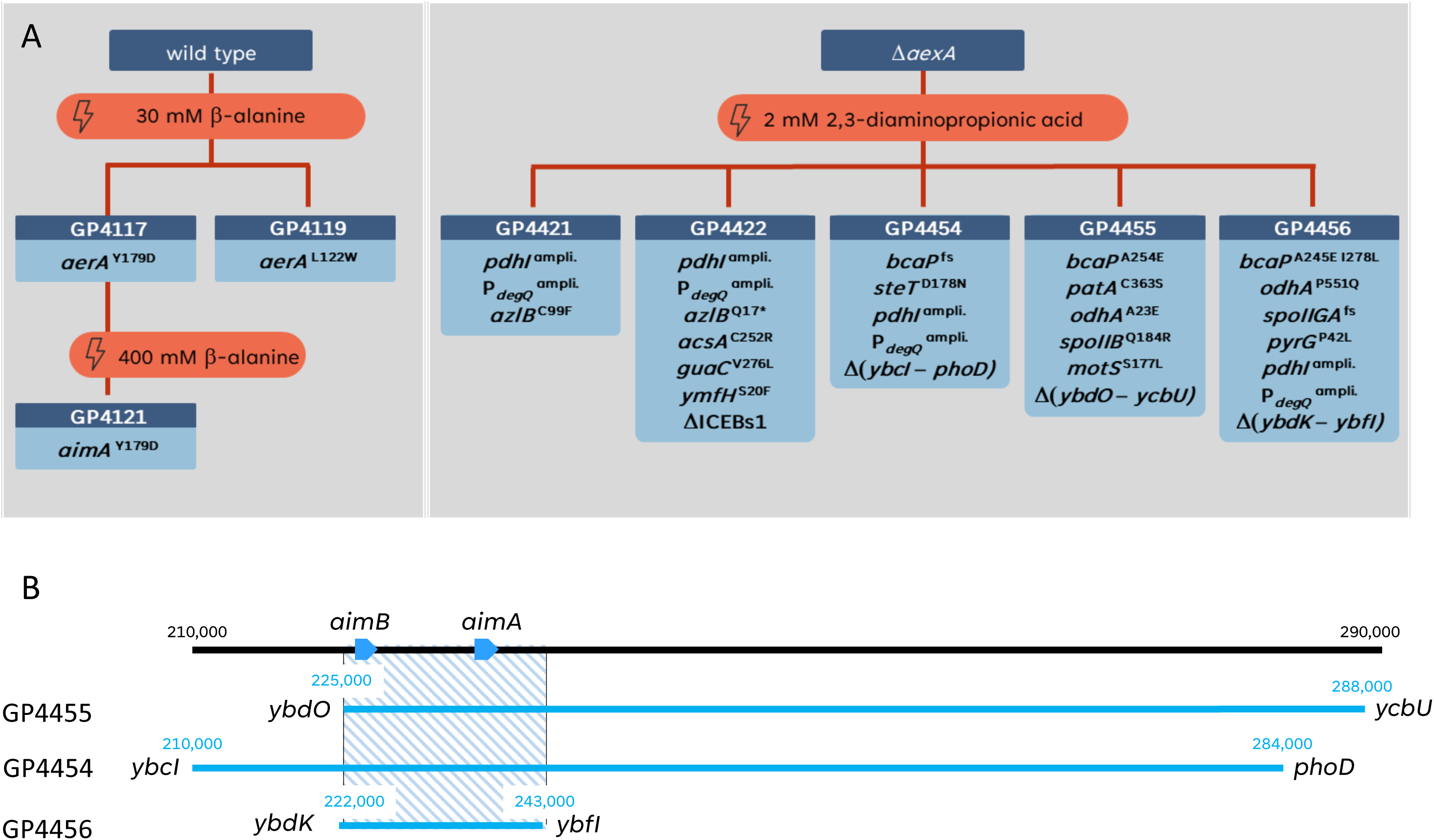
A suppressor screen in the presence of alanine derivates. **A.** Evolutionary trajectory of the wild type strain exposed to toxic concentrations of β-alanine or the Δ*aexA* mutant exposed to toxic concentrations of 2,3-diaminopropionic acid. **B.** Genetic regions affected by deletions in the suppressor strains (GP4454, GP4455, GP4456) under 2,3-diaminopropionic acid stress. Each strain’s deleted regions are demarcated, and an overlay highlights the shared deleted region, denoted by a blue striped rectangle. This shared deleted region suggests a common genetic adaptation in response to the specific stress condition.

Since AerA is a member of the AraC family of transcriptional regulators, the mutations affecting AerA might result in inactivation of the protein or they might have an activating effect. To distinguish between these possibilities, we tested the plating efficiency of the *aerA* deletion mutant BKK05150 on medium containing β-alanine. As shown in Fig. 3A, the mutant was as sensitive as the wild type suggesting that the suppressor mutations resulted in an activation of the AerA protein. In the case of a repressor protein, this would result in even stronger repression of the target genes. In contrast, for an activator, the suppressor mutation might allow transcription activation even in the absence of a putative inducer. As the members of the AraC family are commonly transcription activators (32), it seems most plausible that the latter hypothesis is true. Mutations that result in the activation of otherwise inactive activators are often designated as * mutations. Therefore, we have used the designation *aerA*\* for these suppressor mutations hereafter.

**Figure 3:**
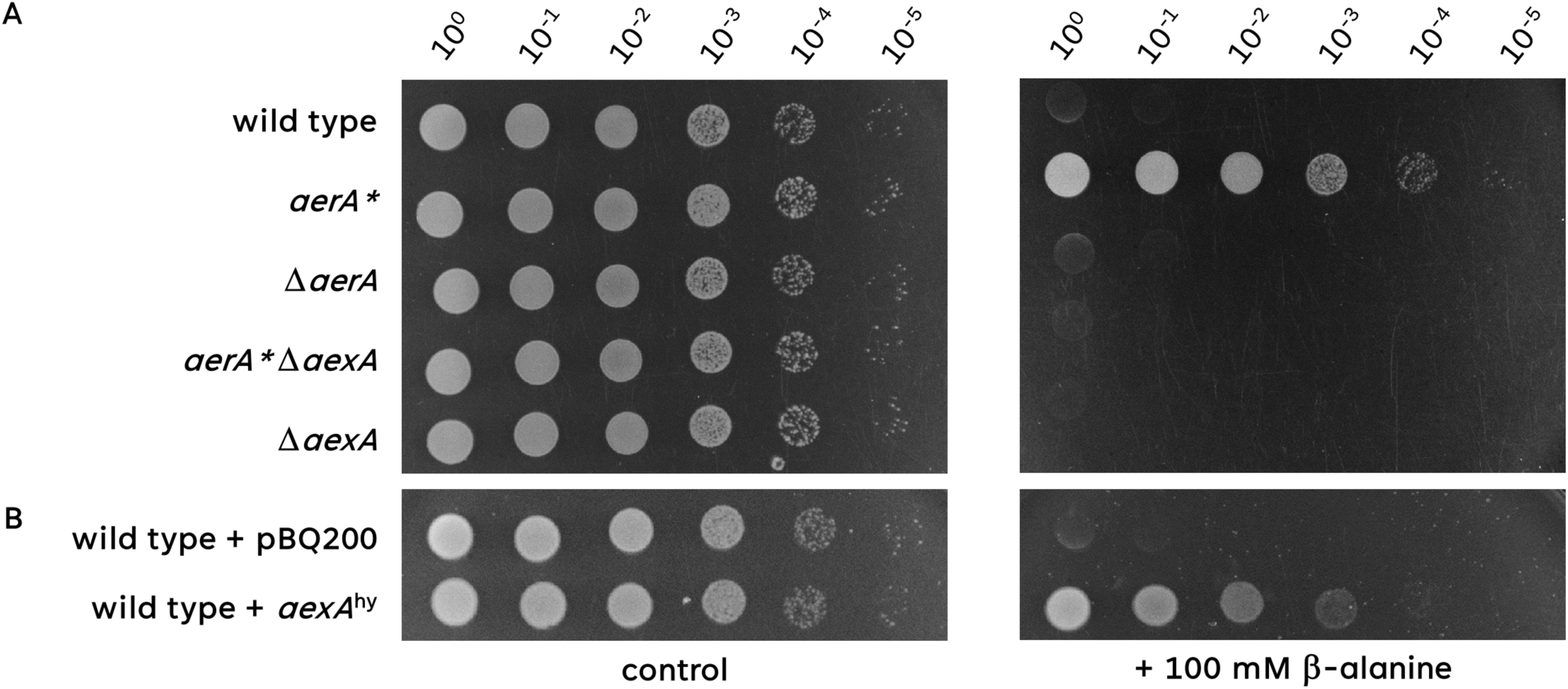
Overexpression of *aexA* is required for β-alanine resistance. **A.** The sensitivity of the wild type, *aerA*\*, Δ*aerA*, *aerA\** Δ*aexA* and Δ*aexA* mutant to β-alanine was tested. The cells were grown in C-Glc minimal medium to an OD_600_ of 1.0 and serial dilutions (10-fold) were prepared. These samples were plated on C-Glc minimal plates containing no β-alanine or 100 mM β-alanine and incubated at 37°C for 48 h. **B.** The sensitivity of the wild type harbouring either the empty vector pBQ200 or pGP3727 (*aexA*^hy^) to β-alanine was tested. The cells were grown in C-Glc minimal medium (with erythromycin and lincomycine) to an OD_600_ of 1.0 and serial dilutions (10-fold) were prepared. These samples were plated on C-Glc minimal plates containing no β-alanine or 100 mM β-alanine and incubated at 37°C for 48 h.

### Overexpression of the EamA family protein AexA is responsible for the tolerance to β-alanine

AraC regulators often control the expression of single genes or operons in their direct genomic vicinity (32). The inspection of the *aerA* genomic context revealed the presence of the *aexA* (*ydeD*) and *ydzE* genes up- and downstream of *aerA*, respectively. The AexA protein is a protein of unknown function that consists of two conserved EamA domains. In the COG database (33), these proteins are annotated as permeases of the drug/metabolite transporter (DMT) superfamily. In contrast, the YdzE gene consists of only one truncated EamA domain, and the Uniprot database suggests that the *ydzE* gene might be a pseudogene. As a putative permease, AexA might be involved in β-alanine homeostasis as an importer or an exporter.

To test whether AexA is involved in β-alanine transport, and whether it acts as an importer or an exporter, we deleted the *aexA* gene in the wild type strain and in the *aerA*\* suppressor mutant and tested the growth of the resulting mutants GP3955 and GP3958 in the presence of β-alanine. Typically, importer mutants show improved growth in the presence of toxic substrates whereas exporter mutants are as sensitive as the parent strain (12, 15, 18, 19). As shown in Fig. 3A, the deletion of *aexA* in the wild type background had no effect on the sensitivity of the bacteria towards β-alanine. In contrast, deletion of *aexA* in the *aerA*\* mutant resulted in the complete loss of the resistance to β-alanine. These observations suggest that AexA most likely acts as an exporter.

If the *aerA*\* mutation would result in the expression of the otherwise silent *aexA* gene, artificial overexpression of the *aexA* gene would be expected to provide tolerance to β-alanine. Therefore, we tested the effect of *aexA* overexpression on the growth in the presence of β-alanine. For this purpose, we used a strain carrying plasmid pGP3727 that allows plasmid-borne constitutive expression of *aexA* in a strain carrying the wild type *aerA* allele. As shown in Fig. 3B, the overexpression of AexA indeed resulted in *aerA*-independent tolerance to β-alanine. Taken together, our findings suggest that AexA is a β-alanine exporter that depends on the mutant AerA* activator protein for expression.

### The *aerA*\* mutation allows inducer-independent activation of the *aexA* promoter

The results presented above suggest that the mutations affecting the AerA protein might result in an inducer-independent activation of AerA that is required for the expression of the otherwise silent *aexA* gene. Previously, the expression of each *B. subtilis* gene has been analyzed under 104 different conditions, including different complex and minimal media (34). The *aexA* gene is barely expressed under all tested conditions (see Fig. 4A). Based on the average expression under all 104 conditions, *aexA* is ranked 3,799^th^ of 4,266 genes. Thus, the condition under which *aexA* is actively expressed has yet to be identified.

**Figure 4:**
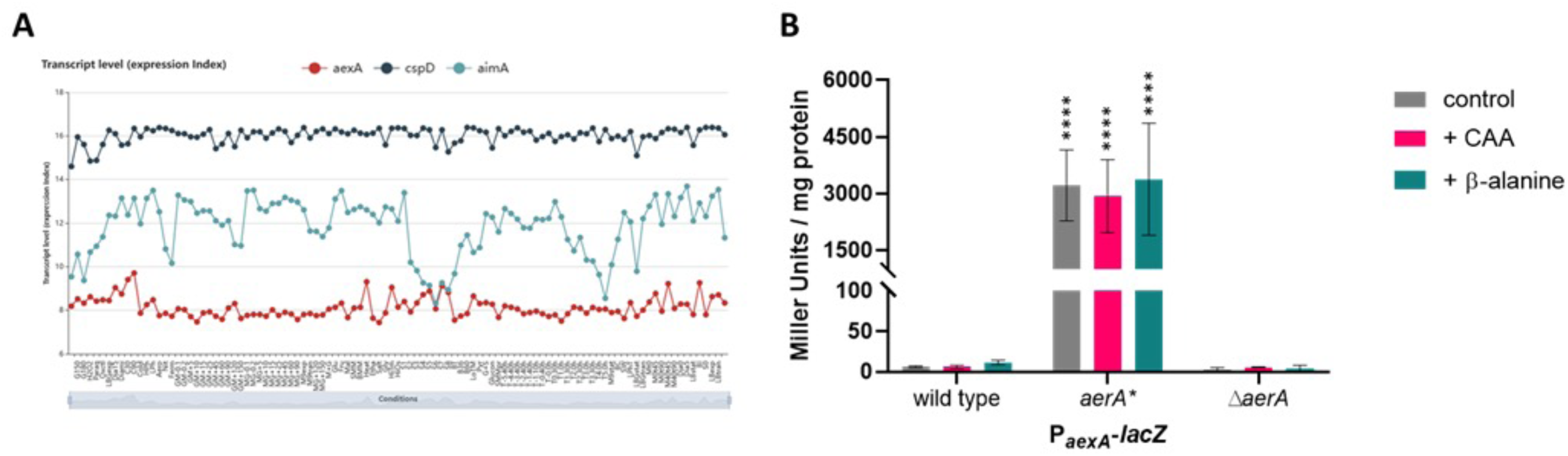
The *aexA* gene is only expressed in the *aerA*\* mutant. **A.** The Expression Browser of SubtiWiki allows a direct comparison of the expression levels of two or more genes (here *aexA, cspD* and *aimA*). The expression level of *cspD* serves as an example of a gene with high constitutive expression. For *aimA*, a moderate expression is observed, while the *aexA* gene is virtually never expressed. **B.** The expression of the *aexA* promotor was monitored in strains that harbor the *aexA*-*lacZ* reporter gene fusion integrated into the chromosomal *amyE* gene. The tested strains were wild type, the *aerA*\* mutant, as well as the deletion mutant Δ*aerA*. Cultures were grown in C-Glc minimal medium supplemented with or without casaminoacids (CAA) (magenta) or β-alanine (cyan) to the early exponential phase (OD_578_ of about 0.6–0.8) and then harvested for β-galactosidase enzyme activity assays. The values for the β-galactosidase activity indicated for each strain represent three independently grown cultures, and for each sample, enzyme activity was determined twice.

To test the role of AerA* in the control of AexA expression, we fused the promoter regions of both the *aerA* and *aexA* genes to a promoterless *lacZ* reporter gene in the background of a wild type strain and the isogenic *aerA*\* and Δ*aerA* mutant strains. The bacteria were grown in C glucose minimal medium and the β-galactosidase activities were determined. Similar results were obtained with the *aexA-lacZ* fusion that reports the activity of the *aexA* promoter (see Fig. 4B). In agreement with the global expression data (Fig. 4A), nearly no expression was observed in the wild type strain GP4132. In contrast, the AerA* protein in GP4438 allowed very high promoter activity (3,210 units per mg of protein) as compared to the expression of genes of central carbon metabolism (35). The deletion of the *aerA* gene in GP4134 resulted in a complete loss of *aexA* expression (Fig. 4B). These data confirm our hypothesis that AerA is a transcription factor that, upon acquisition of an activating * mutation, activates transcription of its own gene and the divergent *aexA* gene. Thus, the suppressor mutation causes tolerance to β-alanine by causing the very high expression of the otherwise silent *aexA* gene.

Often, gene expression can be induced by specific substrates. We therefore also tested the activity of the *aexA* promoter in the three strains during growth in the presence of an amino acid mixture (Casamino acids) and of β-alanine. As shown in Fig. 4B, the addition of β-alanine resulted in a slight increase of promoter activity (6 units per mg of protein to 12 units for the wild type strain GP4132). In contrast, the presence of casamino acids had no effect. Thus, the expression of aexA remained very low even in the presence of β-alanine.

Many regulators control the expression of their own genes. The *aerA* promoter showed little activity in the wild type strain GP4439 (14 units per mg of protein). A significant increase to 42 units was observed in the *aerA*\* mutant GP4440. These data demonstrate that the presence of the AerA* protein results in significant transcriptional activation of its own promoter.

### AexA is a β-alanine exporter

All our genetic data presented above strongly suggest that AexA, if expressed is an exporter for β-alanine. To verify this idea, we investigated the intracellular accumulation of this amino acid by using a radioactively labelled β-alanine. As shown in Fig. 5, both the wild type strain 168 and the isogenic *aerA*\* Δ*aexA* mutant GP3958 accumulated β-alanine in their cells. This observation is also a direct evidence for the existence of uptake system(s) for β-alanine in *B. subtilis*. In contrast, the *aerA*\* mutant GP4117 that exhibits a very high level of *aexA* expression did not accumulate intracellular β-alanine. This is in excellent agreement with the fact that the *aexA* gene is required for β-alanine resistance even in the *aerA*\* mutant (see Fig. 3). Together with the observation that the overexpression of AexA is causative for the tolerance to β-alanine (see above, Fig. 3), the lack of intracellular accumulation of β-alanine upon AexA overexpression clearly demonstrates that AexA indeed acts as an exporter for β-alanine. To the best of our knowledge this is the first identified exporter for a β-amino acid.

**Figure 5:**
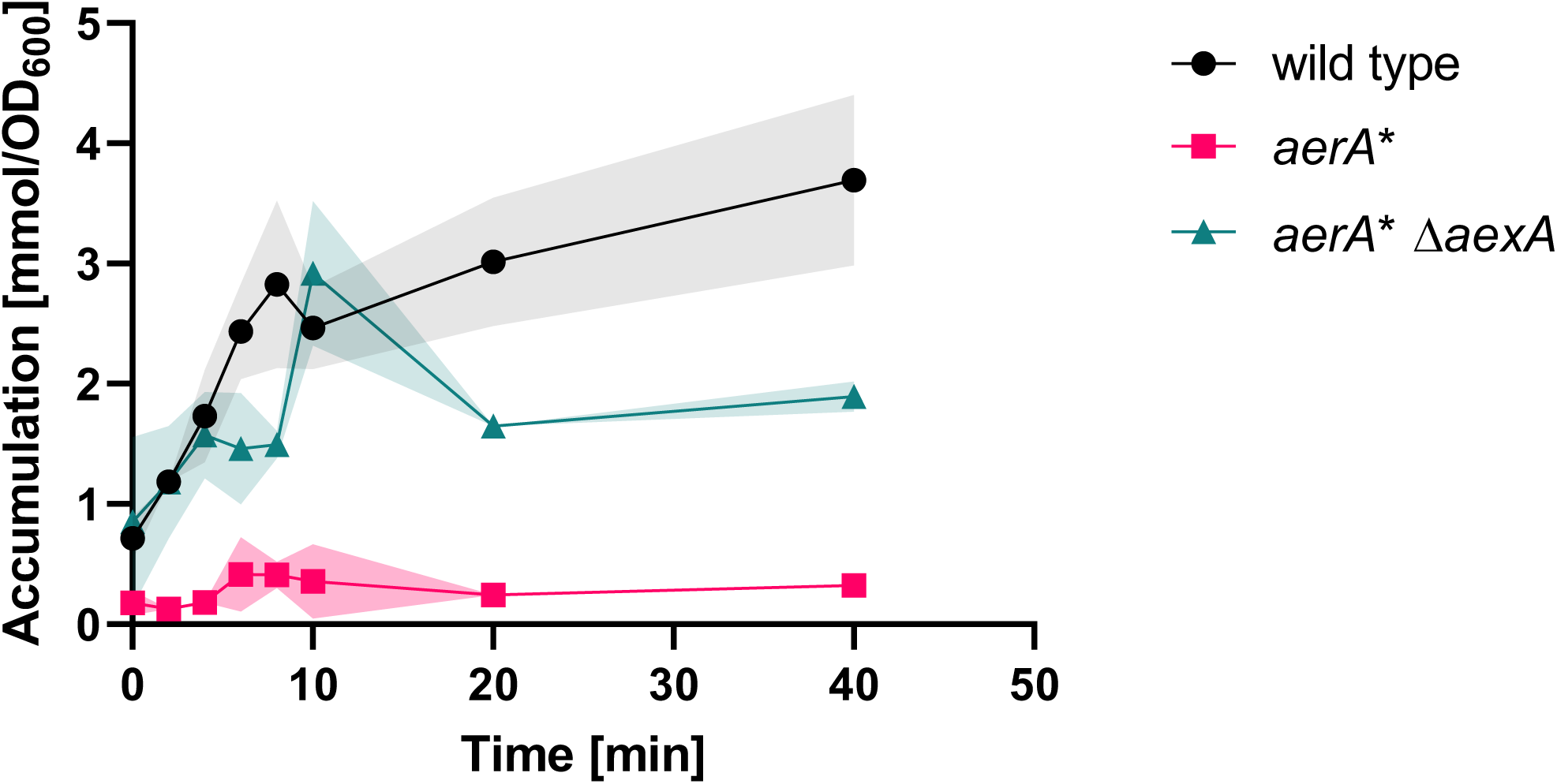
Intracellular accumulation of radiolabeled β-alanine. For the quantification of intracellular accumulation of [1–^14^C]β-alanine, cells of the wild type (black circles), *aerA*\* (magenta squares), and *aerA*\* Δ*aexA* (cyan triangles) mutant were cultivated in C-Glc minimal medium to an OD600 of approximately 0.8. The cells were then diluted with C-Glc minimal medium containing [1–^14^C]β-alanine to reach a final concentration of 30 mM. The accumulation of intracellular [1–^14^C]β-alanine was assayed after 0, 2, 4, 6, 8, 10, 20 and 40 minutes. n=3.

### AexA is a general exporter for pyruvate-based C3-amino acids

Amino acid transporters, both importers and exporters, tend to be promiscuous and can transport multiple amino acids (24). This has been extensively studied for the amino acid importer AimA and the exporter AzlCD which are involved in either the uptake of serine, glutamate, and asparagine or in the export of 4-azaleucine, histidine and asparagine (12, 15, 18, 19, 23). We therefore considered the possibility that AexA might also export amino acids other than β-alanine. As all amino acids that are chemically derived from pyruvate are toxic for *B. subtilis* (12, Fig. 1), we tested the tolerance of a strain overexpressing plasmid-borne AexA to these amino acids, *i.e.* serine, L-alanine and 2,3-diaminopropionic acid. As shown in Fig. 6, AexA expression resulted in increased tolerance to all three amino acids indicating that AexA is also active in the export of these amino acids. The contribution of an *aerA*\* mutation to serine resistance has been described in a recent study (23), and the results reported here demonstrate that this is is caused by the increased expression of AexA in the mutant. We can thus conclude that AexA is a non-specific amino acid exporter for C3-amino acids.

**Figure 6:**
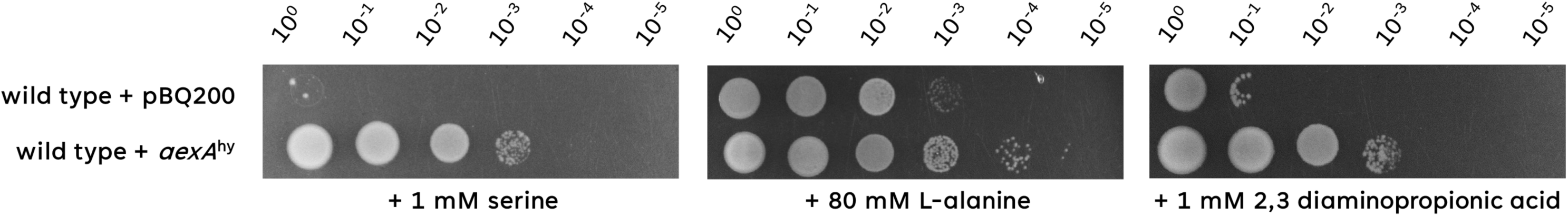
Overexpression of AexA confers resistance against L-serine, L-alanine and 2,3-diaminopropionic acid. The sensitivity of the wild type harbouring the empty vector pBQ200 or pGP3727 *aexA*^hy^ to L-serine, L-alanine or 2,3-diaminopropionic acid was tested. The cells were grown in C-Glc minimal medium to an OD_600_ of 1.0 and serial dilutions (10-fold) were prepared. These samples were plated on C-Glc minimal plates containing 1 mM L-serine, 80 mM L-alanine or 1 mM 2,3-diaminopropionic acid and incubated at 37°C for 48 h.

### 2,3-diaminopropionic acid acts as a weak natural inducer of AerA activity

Of the pyruvate-based amino acids, 2,3-diaminopropionic acid has the most severe growth-inhibiting effect (see Fig. 1). We therefore analyzed the functional link between this amino acid and AerA/AexA in more detail. First, we recorded growth of the wild type strain 168 and the Δ*aexA* mutant GP3955 in the absence and presence of 1 mM 2,3-diaminopropionic acid. As shown in Fig. 7A, the growth of both strains was identical in the absence of the amino acid. In the presence of 2,3-diaminopropionic acid, the wild type strain was able to grow; however, more slowly than in its absence. In contrast, the Δ*aexA* mutant did not grow at all even at this low concentration of 2,3-diaminopropionic acid. This observation confirms the role of AexA in conferring tolerance to 2,3-diaminopropionic acid but also suggests that the AerA is able to provide sufficient activation of AexA expression to allow growth of *B. subtilis* in the presence of 2,3-diaminopropionic acid. To test this idea, we determined the activity of the AerA-dependent *aexA-lacZ* fusion in strain GP4132 in the absence and presence of 2,3-diaminopropionic acid (Fig. 7B). While the *aexA* promoter was inactive in the absence of 2,3-diaminopropionic acid, we observed a 20-fold induction in the presence of the amino acid to 120 units per mg of protein. Even though this promoter activity is much lower than was has been observed with the mutant AerA* protein, it is statistically significant and biologically relevant since this expression is sufficient to provide a basic resistance against 2,3-diaminopropionic acid in *B. subtilis*. Thus, 2,3-diaminopropionic acid might not only be the inducer for AerA, but also a natural substrate for AexA.

**Figure 7:**
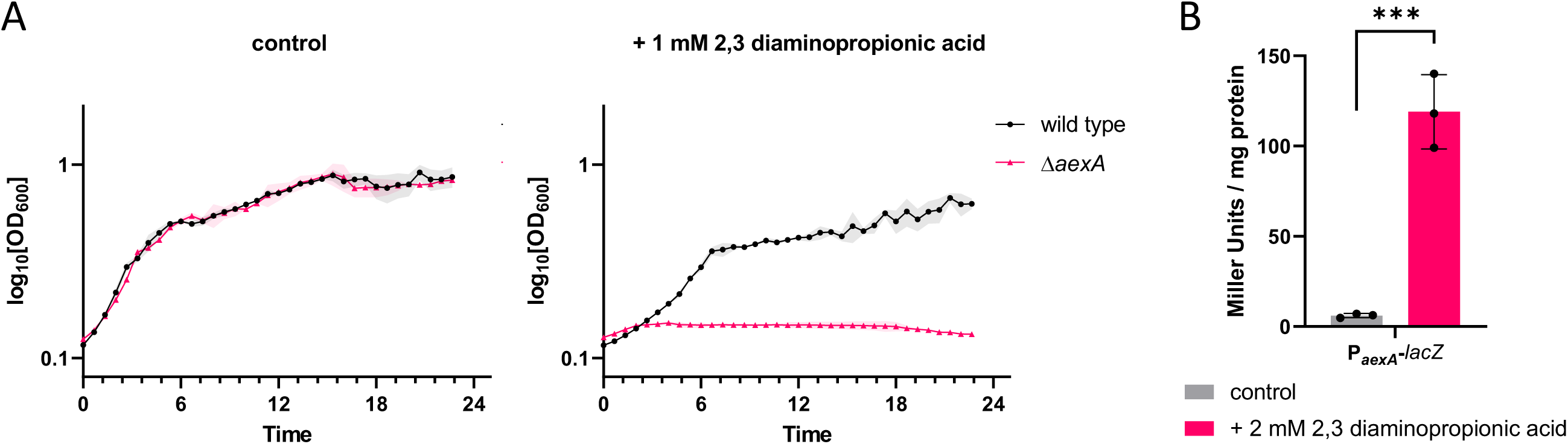
2,3-Diaminopropionic acid is a weak inducer of *aexA* expression. **A.** The growth of the wild type and the Δ*aexA* mutant (GP3955) on C Glc minimal medium was tested in the presence or absence of 1 mM 2,3-diaminopropionic acid. B. The expression of the *aexA* promotor was monitored in strains that harbor the *aexA*-lacZ reporter gene fusion integrated into the chromosomal *amyE* gene. Cultures were grown in C Glc minimal medium in the presence or absence of 2 mM 2,3-diaminopropionic acid to the early exponential phase (OD_578_ of about 0.6–0.8) and then harvested for β-galactosidase enzyme activity assays. The values for the β-galactosidase activity indicated for each strain represent three independently grown cultures, and for each sample, enzyme activity was determined twice.

### Several amino acids protect the cell from the toxic effect of β-alanine

As described above (see Fig. 1), β-alanine is highly toxic in minimal medium, but it is tolerated to high concentrations in complex medium that contains all amino acids. This may indicate that β-alanine interferes with some amino acid biosynthetic pathways, as observed for serine (12). Alternatively, the other, proteinogenic amino acids might compete with β-alanine for uptake capacity thus preventing it from entering the cell. To get further insights into the protection of the cell from toxic β-alanine, we cultivated *B. subtilis* 168 in the presence of β-alanine and each of the proteinigenc amino acids. This analysis revealed that alanine, arginine, isoleucine, and methionine significantly protected the bacteria from β-alanine toxicity (see Fig. 8A). In addition, glycine, proline, and valine had a weak protective effect (see Fig. 8A), whereas the other amino acids did not provide protection. The fact that multiple amino acids protect the cell from β-alanine toxicity suggests that the protection results from competetion for the uptake capacity rather than from an inhibition of specific biosynthetic pathways. We therefore tested the effect of several amino acids on the uptake of β-alanine into the cell. Indeed, we observed a reduced uptake of β-alanine in the presence of alanine, arginine, isoleucine, and methionine, whereas glycine, proline, and valine failed to interfere with β-alanine incorporation (see Fig. 8B). These results correlate well with the results of the plating assay: those amino acids that provide the strongest protection against β-alanine, compete efficiently for the transporter capacity. Thus, β-alanine shares the uptake systems with alanine, arginine, isoleucine and methionine.

**Figure 8:**
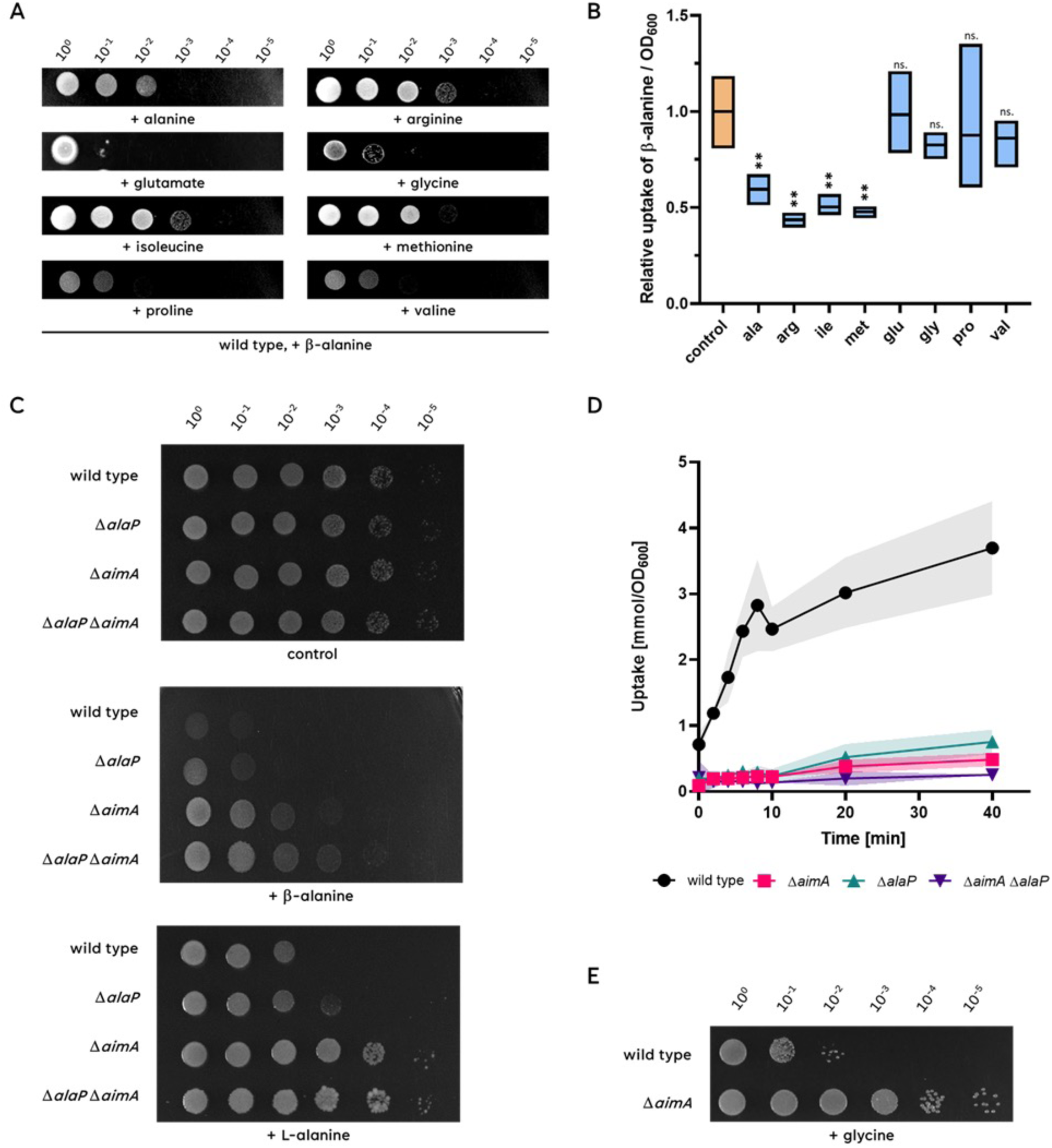
The presence of different amino acids can prevent β-alanine toxicity. **A**. The growth of the wild type strain in presence of β-alanine and either alanine, arginine, isoleucine, methionine, glutamate, glycine, proline or valine was checked. The cells were grown in C-Glc minimal medium to an OD_600_ of 1.0 and serial dilutions (10-fold) were prepared. These samples were plated on C-Glc minimal plates containing 100 mM β-alanine together with 30 mM of the respective amino acid and incubated at 37°C for 48 h. **B.** To check the relative uptake of β-alanine in presence or absence of the tested amino acid, the wild type was grown in C-Glc minimal medium containing 30 mM β-alanine spiked with radiolabeled [1–^14^C]β-alanine with or without 5 mM of the respective amino acid to an OD_600_ of approximately 0.8. The accumulation of intracellular [1–^14^C]β-alanine was assayed after 40 minutes. n=3 **C.** The sensitivity of the wild type 168, Δ*alaP* (GP4136), Δ*aimA* (GP3590)*, and* Δ*alaP* Δ*aimA* (GP4135) to β-alanine or L-alanine was tested. The cells were grown in C-Glc minimal medium to an OD_600_ of 1.0 and serial dilutions (10-fold) were prepared. These samples were plated on C-Glc minimal plates containing 100 mM β-alanine or L-alanine and incubated at 37°C for 48 h. **D.** For the quantification of intracellular accumulation of [1 – 14 C] β-alanine, cells of the wild type (168, black circles), Δ*alaP* (GP4136, cyan triangles pointing upwards), Δ*aimA* (GP3590, magenta squares)*, and* Δ*alaP* Δ*aimA* (GP4135, purple triangles pointing downwards) mutant were cultivated in C-Glc minimal medium to an OD600 of approximately 0.8. The cells were then diluted with C-Glc minimal medium containing [1–^14^C]β-alanine to reach a final concentration of 30 mM. The accumulation of intracellular [1–^14^C]β-alanine was assayed after 0, 2, 4, 6, 8, 10, 20 and 40 minutes. n=3 **E.** The sensitivity of the wild type 168 and the Δ*aimA* (GP3590), to glycine was tested. The cells were grown in C-Glc minimal medium to an OD_600_ of 1.0 and serial dilutions (10-fold) were prepared. These samples were plated on C-Glc minimal plates containing 200 mM glycine and incubated at 37°C for 48 h.

### AlaP and AimA are the importers for both L-α-alanine and β-alanine

As described above, β-alanine and alanine seem to share their transporter(s). A recent study has identified the AlaP protein as the transporter for D-α-alanine (36). Based on the structural similarity of both amino acids and on the observed competition for uptake capacity between them (see Fig. 8B), we hypothesized that AlaP might also be involved in the transport of β-alanine. Moreover, our suppressor screen already identified a mutation in the low affinity amino acid importer AimA that results in resistance to very high concentrations of β-alanine that are not even tolerated if the amino acid exporter AexA is expressed due to the *aerA*\* mutation (GP4121, see Fig. 2). This obervation suggests that AimA also imports β-alanine in addition to serine, glutamate and aspartate (12, 15, 23).

To test the possible involvement of AlaP and AimA in the transport of both amino acids, we tested the growth of a wild type strain and single and double mutants, *alaP* and *aimA*, in the presence of otherwise toxic concentrations of both amino acids. As shown in Fig. 8C, the *alaP* mutant GP4146 has a slightly higher tolerance towards both β-alanine and L-alanine than the wild type strain, indicating that AlaP contributes to the uptake of both amino acids. The *aimA* mutant GP3590 exhibited tolerance towards both amino acids, even more pronounced as compared to the *alaP* mutant. While the lack of AimA did not provide full resistance to β-alanine as observed for the *aerA*\* mutant, the *aimA* mutant fully tolerated the presence of L-alanine. Thus, our results suggest that both AlaP and AimA contribute to the uptake of both, β-alanine and L-alanine. Indeed, the double mutant GP4135 showed increased resistance to both amino acids as compared to the two isogenic single mutants (see Fig. 8C). This confirms the idea, that both AlaP and AimA contribute to the uptake of β-alanine and L-alanine.

To provide direct evidence for β-alanine uptake by AlaP and AimA, we measured the β-alanine transport of a wild type strain (*B.subtilis* 168), an *alaP* mutant (GP4146), an *aimA* mutant (GP3590) and a *alaP aimA* double mutant (GP4135). As shown in Fig. 8D, β-alanine was efficiently transported by the wild type strain, whereas a severe reduction was observed in the *alaP* and *aimA* mutants. In the *alaP aimA* double mutant GP4135, the transport of β-alanine was nearly completely abolished (see Fig. 8D), confirming that AlaP and AimA are the main players in β-alanine uptake in *B. subtilis*.

### AimA is responsible for the uptake of glycine

With the results presented above, amino acids importers have been identified in *B. subtilis* for all proteinogenic amino acids with the exception of glycine, phenylalanine and tyrosine. As AimA transports all so far tested C3 amino acids, we considered the possibility that it might also be involved in the uptake of the C2 amino acid glycine (see Fig. 8E). As observed for alanine, glycine also inhibited growth of *B. subtilis* 168 on minimal medium. In contrast, the isogenic *aimA* mutant GP3590 was resistant to this amino acid indicating that AimA is indeed the main glycine permease in B. subtilis.

### AimA, AimB, and BcaP are the importers for 2,3-diaminopropionic acid

To get first insights into the uptake of 2,3-diaminopropionic acid, we isolated suppressor mutants that are able to grow in the presence of 2 mM of this amino acid. Since the exporter AexA provides already a weak natural resistance towards diaminopropionic acid, we used the *aexA* mutant GP3955 for suppressor isolation to ensure that we identify AexA-independent players in diaminopropionate homeostasis. The determination of the genome sequences of the five independent suppressor mutants revealed the presence of multiple mutations in each mutant (see Fig. 2A). Interestingly, three of the mutants had genomic deletions of 21 to 74 kb. In all these mutants the deletion affected the region from *ybdO* to *ybfI* as the common denominator (see Fig. 2B). This region encodes two known amino acid transporters, AimA and AimB (previously YbxG). Moreover, these three strains had mutations affecting the amino acid transporter BcaP, in one case a frameshift mutation that results in the synthesis of a truncated inactive protein. To study the role of these three amino acid transporters in providing resistance to 2,3-diaminopropionic acid, we tested the growth of the corresponding single, double, and triple mutants in the presence of this amino acid. As shown in Fig. 9A, the single mutants were still strongly inhibited by 2,3-diaminopropionic acid. The double mutants showed increased resistance which was most pronounced in the mutants lacking AimA in addition to either BcaP or AimB. The triple Δ*aimA* Δ*bcaP* Δ*aimB* mutant GP4495 had the highest resistance, with growth only weakly affected. These observations demonstrate that all three transporters, AimA, BcaP, and AimB, transport 2,3-diaminopropionic acid into the cell, thus inactivation of all three as observed in all suppressor mutants is required for a high-level resistance of *B. subtilis* to 2,3-diaminopropionic acid. The identification of diaminopropionate as an additional substrate for AimB suggests that this protein also is a general broad-spectrum amino acid permease.

**Figure 9:**
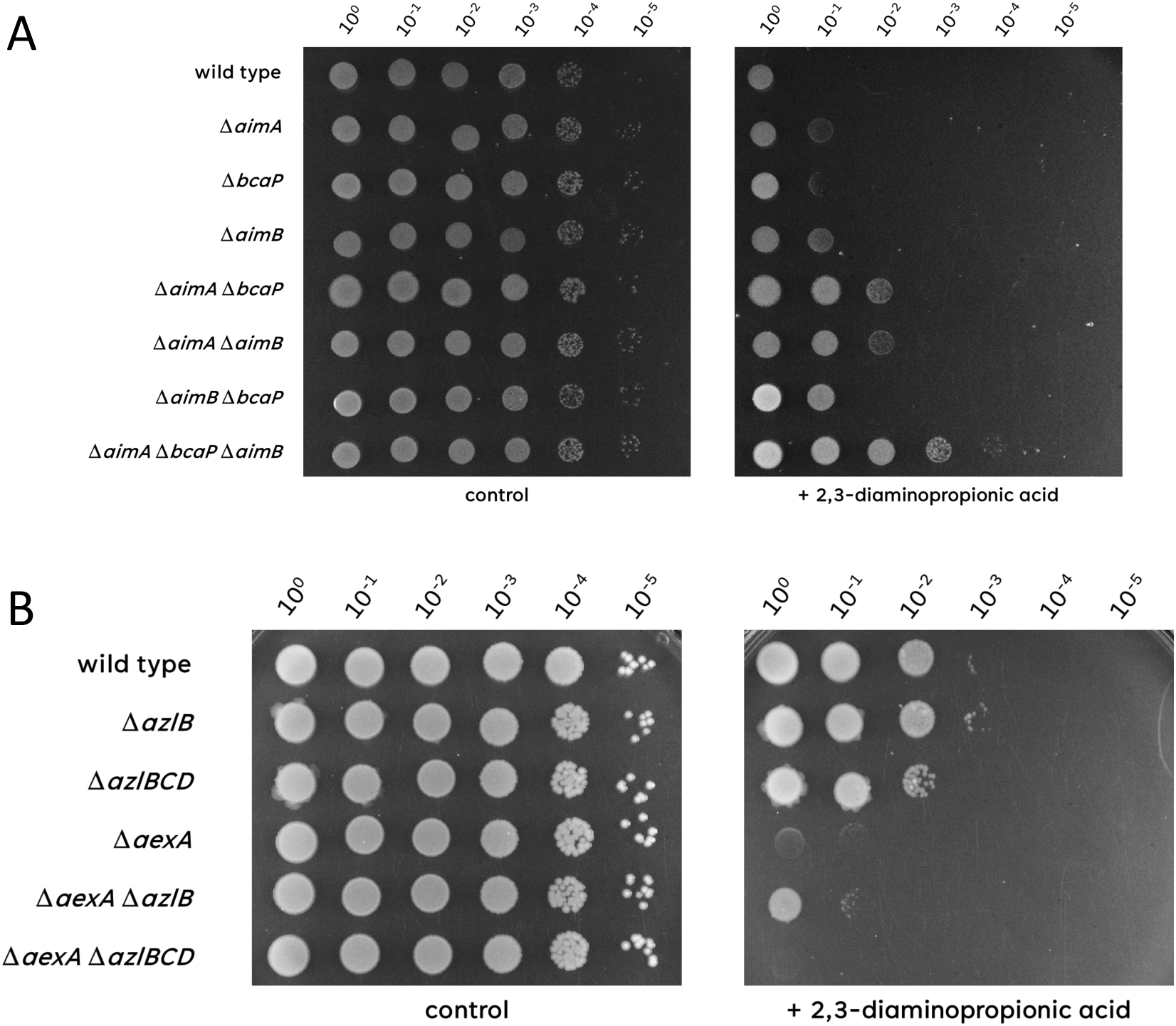
Sensitivity of mutant strains to 2,3-diaminiopropionic acid. **A.** The sensitivity of the wild type (168), Δ*aimA* (GP2786), Δ*bcaP* (GP4484), Δ*aimB* (GP2396), Δ*aimA* Δ*bcaP* (GP2949), Δ*aimA* Δ*aimB* (GP2951), Δ*aimB* Δ*bcaP* (GP2952)*, and* Δ*aimA* Δ*aimB* Δ*bcaP* (GP2950) to 2,3-diaminopropionic acid was tested. The cells were grown in C-Glc minimal medium to an OD_600_ of 1.0 and serial dilutions (10-fold) were prepared. These samples were plated on C-Glc minimal plates containing no, 100 mM, or 200 mM β-alanine and incubated at 37°C for 48 h. **B.** The sensitivity of the wild type (168), Δ*azlB* (GP3600), Δ*azlB*CD (GP3623), Δ*aexA* (GP3955), Δ*aexA* Δ*azlB* (GP4384), and Δ*aexA* ΔazlBCD (GP4385) to 2,3-diaminopropionic acid was tested. The cells were grown in C-Glc minimal medium to an OD_600_ of 1.0 and serial dilutions (10-fold) were prepared. These samples were plated on C-Glc minimal plates containing no, 100 mM, or 200 mM β-alanine and incubated at 37°C for 48 h.

The remaining two 2,3-diaminopropionic acid-tolerant mutants both had mutations affecting the transcription repressor AzlB that controls the expression of the well-studied amino acid exporter AzlCD (18, 19, 23) (see Fig. 2). The *azlB* mutations result in high expression of the otherwise poorly expressed broad-spectrum amino acid exporter AzlCD. To test the role of AzlB and the AzlCD amino acid transporter in conferring resistance to 2,3-diaminopropionic acid, we compared the growth of the *azlB* and *azlBCD* mutants in the genetic backgrounds of a wild type strain and of an *aexA* mutant that is unable to export 2,3-diaminopropionic acid via AexA. (Fig. 9B). The latter mutant had been used to isolate the resistant mutants (see Fig. 2). In the wild type background, the Azl system had no effect on the resistance towards 2,3-diaminopropionic acid, probably due to the presence of the natural exporter AexA. Indeed, an *azlB* mutant only conferred increased 2,3-diaminopropionic tolerance to the *aexA* mutant and this tolerance depends on the presence of the *azlCD* exporter genes indicating that the amino acid exporter AzlCD can contribute to 2,3-diaminopropionic acid export in addition to AexA.

## DISCUSSION

The results presented in this study identify the transport systems for C2 and C3 amino acids in *B. subtilis* (see Fig. 10). All these amino acids can become toxic for *B. subtilis* if they are present at high concentrations. Thus, mutant characterization and suppressor screens are excellent tools to get insights into both the active and dormant transport systems.

**Figure 10:**
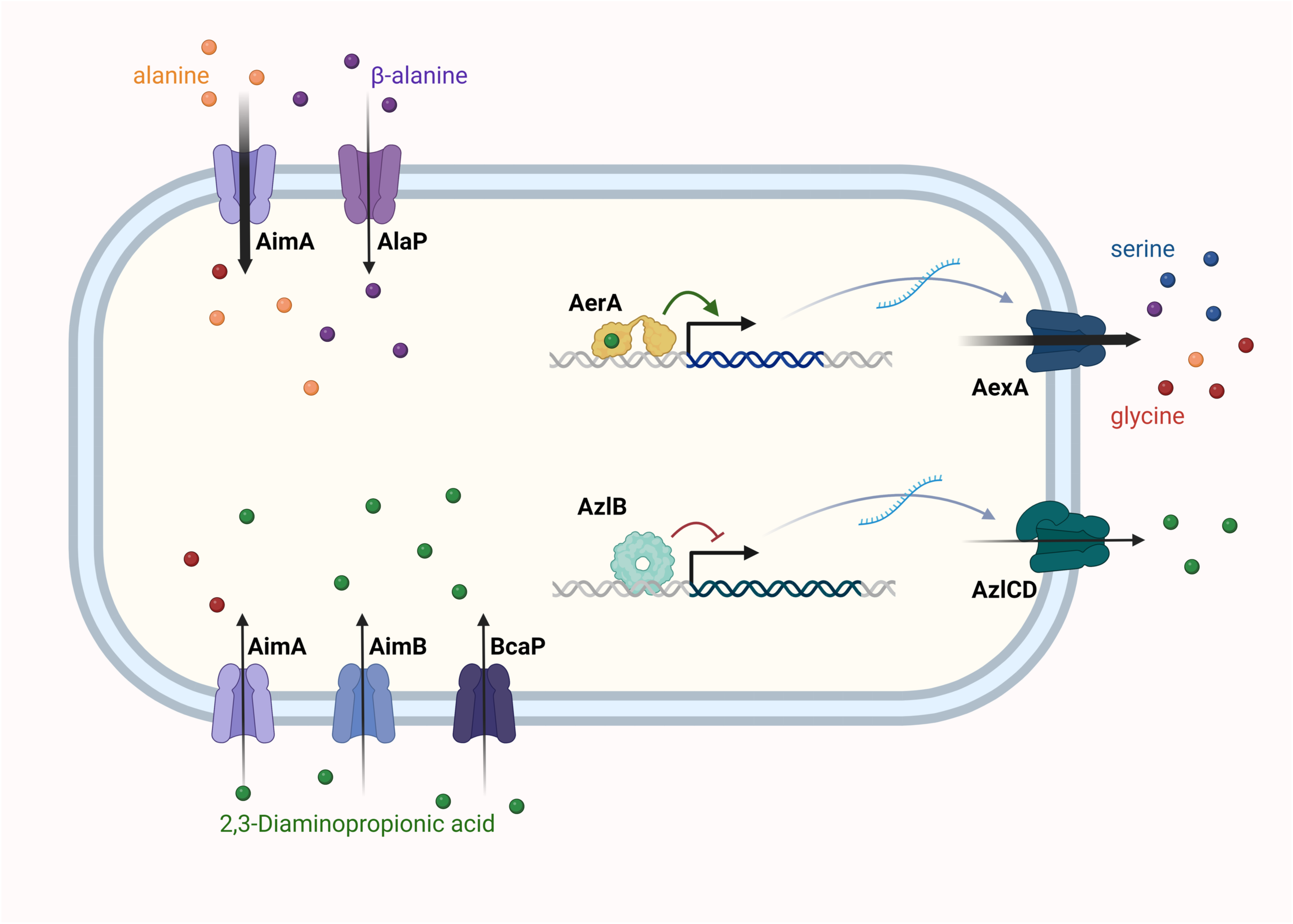
Transport of C2 and C3 amino acids in *B. subtilis*. The uptake and export systems for the C2 and C3 amino acids are shown in the left and right part of the figure, respectively. The amino acid exporters AexA and AzlCD are only expressed if the transcription factors AerA and AzlB have acquired mutations that allow their DNA binding in the absence of the inducer or in the presence of 2,3-diaminopropionic acid (for AerA).

### Broad-range APC family amino acid permeases can transport multiple amino acids

Our work identifies AimA as a major broad-range amino acid transporter in this *B. subtilis*. This permease is able to take up glycine, L-alanine, β-alanine, and 2,3-diaminopropionic acid in addition to glutamate, asparagine, and serine (12, 15, 23). This role is in good agreement with the constitutive expression of the *aimA* gene under a broad variety of conditions (see Fig. 4A) (34). As already observed for several other amino acids, the C3 amino acids are all transported by more than one permeases. In the case of L-alanine and β-alanine, this is AlaP which was also identified as a transporter for D-alanine (36). Surprisingly, 2,3-diaminopropionic acid can even be transported into the cell by three amino acid permeases, the promiscuous transporters AimB (previously YbxG) and BcaP in addition to AimA. These transporters have previously been shown to be required for the uptake of threonine and serine (12, 20, 37). In addition, BcaP also acts in the acquisition of asparagine and the branched-chain amino acids isoleucine and valine (23, 37). It is tempting to speculate, that 2,3-diaminopropionic acid as a non-natural growth-inhibiting amino acid can be transported by multiple transporters, as no selectivity against this molecule is required in natural ecosystems. Interestingly, all four of these amino acid permeases involved in the uptake of alanine, β-alanine and 2,3-diaminopropionic acid are members of the large amino acid-polyamine-organocation (APC) superfamily (38). These proteins typically transport amino acids, but they may also be involved in the uptake of unrelated substrates such as methyl-thioribose (MtrA) or potassium ions (KimA) (39, 40).

The APC family transporters are ubiquitous in all domains of life with typically between 20 to 30 members in each organism. Poor specificity seems to be a rather general characteristic of this family, suggesting that they are important for the acquisition of a large range of different amino acids and other compounds in archaea, bacteria, yeast and man. However, the individual melange of these proteins differs from species to species. Interestingly, AimA seems to be limited to *B. subtilis*, as close orthologs of the protein are missing in most species, even in other *Bacillus* species.

### The power of genetic screens: You get what you select for

It is interesting to note that three of the suppressor mutants that were resistant to 2,3-diaminopropionic acid contained large genomic deletions. In each case, the deletions resulted in the loss of AimA and AimB (see Fig. 2B). In two cases, GltP, a third putative amino acid permease that was shown to be involved in the transport of the antimetabolite glyphosate (22), was also missing due to the deletions. However, in GP4456, which had the smallest deletion, the *gltP* gene was still present suggesting that it was not involved in the sensitivity to 2,3-diaminopropionic acid. In prior studies, numerous *aimA* mutants that were resistant to either serine, glutamate, or asparagine have been isolated. However, all of them just had single nucleotide polymorphisms or small deletions or insertions within the *aimA* gene but not deletions of complete genomic regions (12, 15, 23). The ubiquitous occurrence of deletions of the *aimA-aimB* genomic region indicates that the selective pressure was really directed to the loss of both transporters (in addition to BcaP) and underlines the power of genetic screens for the identification of genes and proteins that are involved in so far poorly studied functions.

### Mutational activation of otherwise cryptic genes is a common theme in the adaptation to toxic amino acids

Previous studies have identified the AzlCD amino acid exporter to be required for the disposal of harmful amino acids in the cases of 4-azaleucine, histidine and asparagine in *B. subtilis* and this study indicates that AzlCD overexpression also contributes to the resistance to 2,3-diaminopropionic acid (18, 19, 23, 41). In each case, expression of the AzlCD exporter and subsequent resistance to these amino acids was acquired by the mutational inactivation of the AzlB repressor either by amino acid substitutions or by frame-shift mutations. Similarly, mutations in the AerA transcription activation trigger the expression of the AexA amino acid exporter. In this case, only point mutations that allow DNA binding and subsequent transcription activation in the absence of any cofactor are possible. Similar mutations that render transcription activators effector-independent have been identified for the Crp and PrfA regulators in *E. coli* and *Listeria monocytogenes* that control the expression of genes for the utilization of secondary carbon sources and of virulence factors, respectively (42, 43, 44). Interestingly, in a recent study aimed at the identification of *B. subtilis* genes and proteins involved in the sensitivity to asparagine, a *yetL* mutant was observed (23). Strikingly, YetL is a MarR-type transcription factor that is encoded in the genome next to YetK, another EAM family transporter. It is tempting to speculate that the mutation in resulted again in effector-independent activity of YetL.

### A strongly conserved family of poorly expressed “sleeping beauty amino acid exporters”

As mentioned above, AexA is a member of the EamA transporter family, named after the *E. coli* cysteine/O-acetylserine exporter EamA (45). These proteins consist of two conserved EamA domains (46). According to the COG database (33), members of this family are ubiquitous in all organisms, with multiple proteins in most species. Characterized members of the family include EamA and YddG from *E. coli*, which export cysteine/O-acetylserine and aromatic amino acids, respectively (45, 47). *B. subtilis* encodes eight members of the EamA transporter family. One particularly striking common feature of the EamA type transporter genes in *B. subtilis* is their low expression under all studied conditions (see Fig. 4A with *aexA* as a representative example, 34). Most of the corresponding genes belong to the 10% of the most poorly expressed genes in *B. subtilis*, and the *ydfC* gene encoding one of these uncharacterized transporters has the third position in the list of the most poorly expressed genes in *B. subtilis*. It is possible that the conditions that induce the expression of the EamA transporters have not been identified, as is the case for AexA which is induced in the presence of 2,3-diaminopropionic acid. Alternatively, it is possible they are never expressed unless the cells experience a selective pressure, that results in the acquisition of mutations that allow expression of the transporters, such as the presence of a toxic substrate. Another example is the EAM family member YetK which may become expressed upon the acquisition of YetL* mutations in response to asparagine stress (23, see above). It is tempting to speculate that YetK is active in asparagine export. Based on amino acid export as the common function of these transporters and their commonly very low and only occasionally induced expression, we suggest to dub this protein family “sleeping beauty amino acid exporters”.

With the identification of transporters for the C2 and C3 amino acids in *B. subtilis*, phenylalanine and tyrosine remain the last amino acids, for which no permease has so far been identified in this bacterium. While the field of amino acid uptake is thus coming to an end, the export of amino acid still remains an interesting area of research. It will be interesting how *B. subtilis* responds to the presence of other non-natural amino acids, and whether the activation of sleeping beauty amino acid exporters helps in the adaptation to such molecules.

## MATERIALS AND METHODS

### Strains, media and growth conditions

*E. coli* DH5α (48) was used for cloning. All *B. subtilis* strains used in this study are derivatives of the laboratory strain 168. They are listed in Table 1. *B. subtilis* and *E. coli* were grown in Luria-Bertani (LB) or in sporulation (SP) medium (48, 49). For growth assays, *B. subtilis* was cultivated in C minimal medium containing 0.5% glucose (50) or MSSM medium (51). MSSM is a modified SM medium in which KH_2_PO_4_ was replaced by NaH_2_PO_4_ and KCl was added as indicated (51). The media were supplemented with ampicillin (100 µg/ml), kanamycin (10 µg/ml), chloramphenicol (5 µg/ml), spectinomycin (150 µg/ml), or erythromycin and lincomycin (2 and 25 µg/ml, respectively) if required.

**Table 1.**
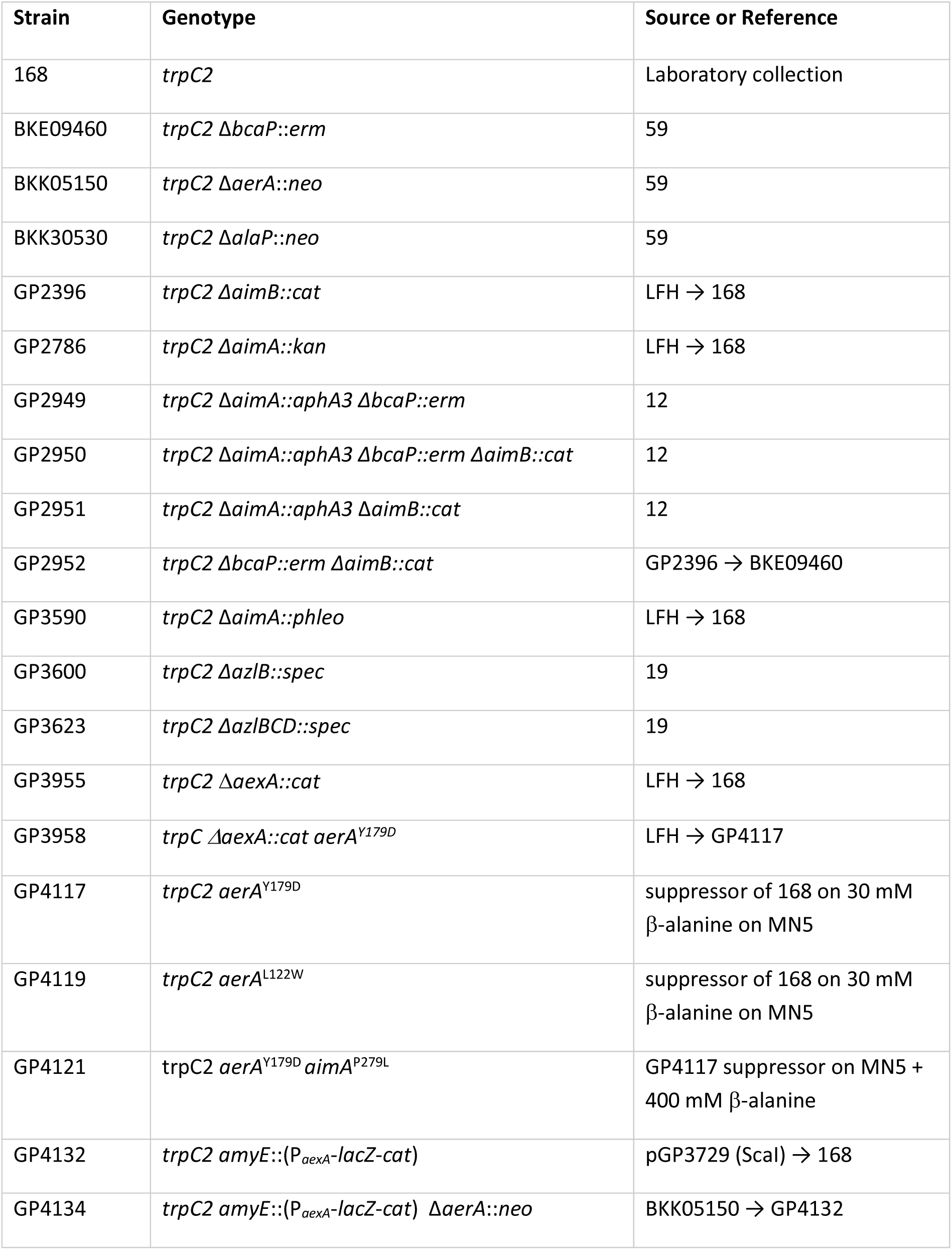
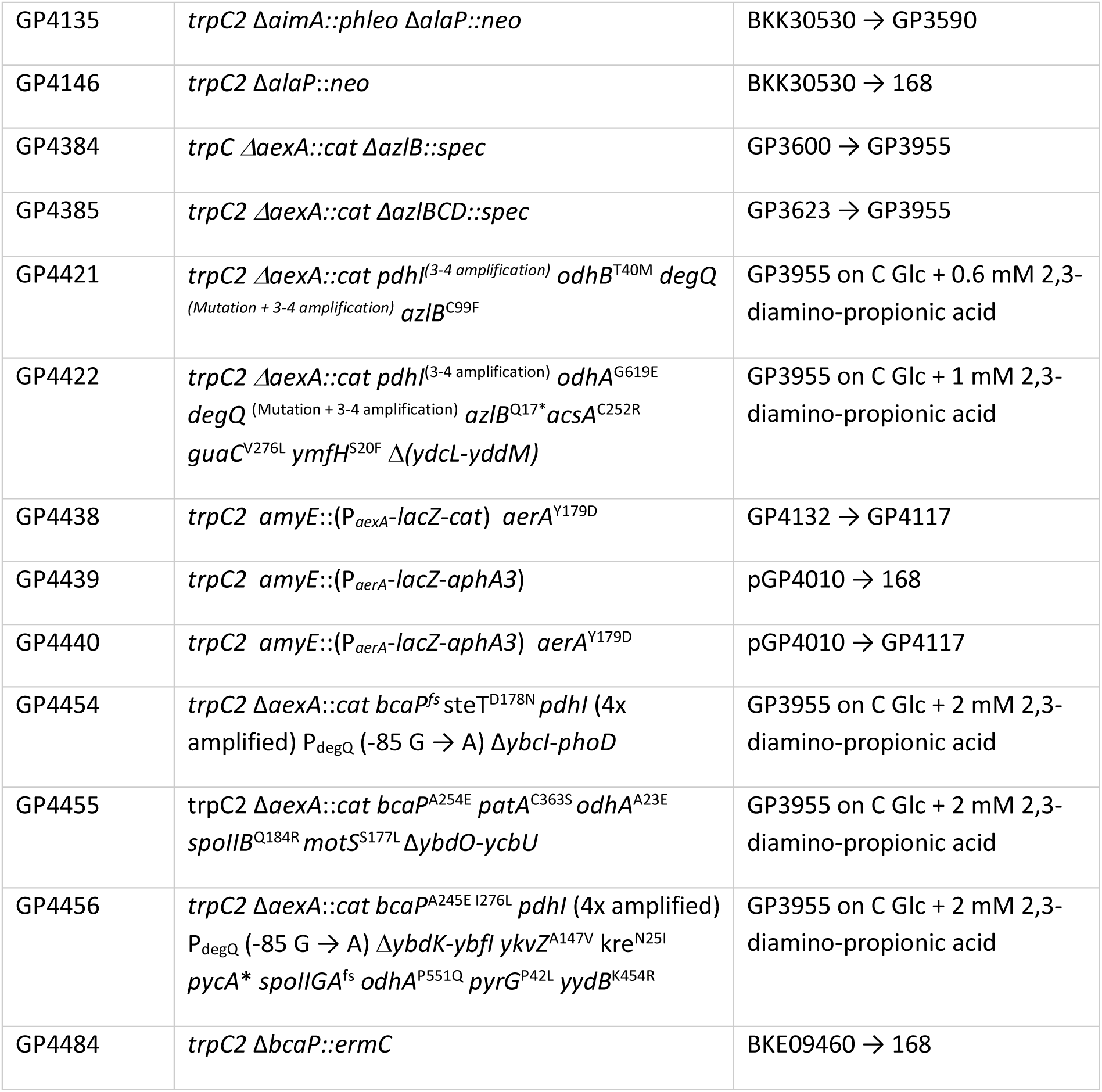
*B. subtilis* strains used in this study.

### DNA manipulation and transformation

Transformation of *E. coli* and plasmid DNA extraction were performed using standard procedures (48). All commercially available plasmids, restriction enzymes, T4 DNA ligase and DNA polymerases were used as recommended by the manufacturers. *B. subtilis* was transformed with plasmids, genomic DNA or PCR products according to the two-step protocol (49). Transformants were selected on LB plates containing antibiotics suitable for the introduced construct. DNA fragments were purified using the QIAquick PCR Purification Kit (Qiagen, Hilden, Germany). DNA sequences were determined by the dideoxy chain termination method (48).

### Construction of mutant strains by allelic replacement

Deletion of the *aimA*, *aimB* and *aexA* genes was achieved by transformation of *B. subtilis* 168 with a PCR product constructed using oligonucleotides to amplify DNA fragments flanking the target genes and an appropriate intervening resistance cassette as described previously (52). The integrity of the regions flanking the integrated resistance cassette was verified by sequencing PCR products of about 1,100 bp amplified from chromosomal DNA of the resulting mutant strains (see Table 1).

### Phenotypic analysis

In *B. subtilis*, amylase activity was detected after growth on plates containing nutrient broth (7.5 g/l), 17 g Bacto agar/l (Difco) and 5 g hydrolyzed starch/l (Connaught). Starch degradation was detected by sublimating iodine onto the plates.

Quantitative studies of *lacZ* expression in *B. subtilis* were performed as follows: cells were grown in C-glucose minimal medium. Cells were harvested at OD_600_ of 0.5 to 0.8. β-Galactosidase specific activities were determined with cell extracts obtained by lysozyme treatment as described previously (49). One unit of β-galactosidase is defined as the amount of enzyme which produces 1 nmol of o-nitrophenol per min at 28° C.

### Genome sequencing

To identify the mutations in the suppressor mutant strains (see Fig. 2 and Table 1), the genomic DNA was subjected to whole-genome sequencing. Concentration and purity of the isolated DNA was first checked with a Nanodrop ND-1000 (PeqLab Erlangen, Germany), and the precise concentration was determined using the Qubit® dsDNA HS Assay Kit as recommended by the manufacturer (Life Technologies GmbH, Darmstadt, Germany). Illumina shotgun libraries were prepared using the Nextera XT DNA Sample Preparation Kit and subsequently sequenced on a MiSeq system with the reagent kit v3 with 600 cycles (Illumina, San Diego, CA, USA) as recommended by the manufacturer. The reads were mapped on the reference genome of *B. subtilis* 168 (GenBank accession number: NC_000964) (53). Mapping of the reads was performed using the Geneious software package (Biomatters Ltd., New Zealand) (54). Frequently occurring hitchhiker mutations (55) and silent mutations were omitted from the screen.

The resulting genome sequences were compared to that of our in-house wild type strain. Single nucleotide polymorphisms were considered as significant when the total coverage depth exceeded 25 reads with a variant frequency of ≥90%. All identified mutations were verified by PCR amplification and Sanger sequencing.

### Plasmid constructions

The plasmids pAC6 and pAC7 (56, 57) were used to construct translational fusions of the potential *aexA* and *aerA* promoter regions, respectively, to the promoterless *lacZ* gene. For this purpose, the promoter regions were amplified using oligonucleotides that attached EcoRI and BamHI restriction to the ends of the products. The fragments were cloned between the EcoRI and BamHI sites of the plasmids. The resulting plasmids were pGP3729 (*aexA-lacZ*) and pGP4010 (*aerA-lacZ*).

To allow for the ectopic expression of the *aexA* gene, we constructed the plasmid pGP3727, respectively. The corresponding gene were amplified using oligonucleotides that added SphI and SalI sites to the ends of the fragments and cloned into the replicative expression vector pBQ200 (58) that was linearized with the same enzymes.

### Intracellular accumulation of radiolabeled β-alanine in *B. subtilis*

To study the intracellular accumulation of [1–^14^C]β-alanine in cells of the *B. subtilis*, cultures were grown in C minimal medium containing glucose to exponential growth phase (OD_600_, 0.7-0.9) and β-alanine was spiked in with radiolabeled [1–^14^C]β-alanine (specific activity for both solutes, 50-60 mCi/mmol) was added to a final concentration of 30 mM. Uptake of the amino acid was followed at 0, 2, 4, 6, 8, 10, 20 and 40 minutes by pipetting 500 µl of the culture on a filter. The cells were washed 3 times with 1 ml of C minimal medium and the filter was transferred to a scintillation vial. The filters were completely dried for 30 min at 70°C, before adding 5 ml Ecoscint A liquid scintillation cocktail (National Diagnostics, USA) was added and the radioactivity was measured using a HIDEX 300SL β-particle scintillation detector. For the controls, 500 µl of C Glc with 30 mM β-alanine with and without radiolabeled [1–^14^C]β-alanine were pipetted directly into the scintillation vial and let dried completely at 70°C.

### Competitive uptake of radiolabeled β-alanine in *B. subtilis* in the presence of amino acids

The competitive uptake of β-alanine in the presence of other amino acids L-alanine, L-arginine, L-isoleucine, L-methionine, L-glutamate, glycine, L-proline and L-valine was measured. The *B. subtilis* wild type strain 168 was cultivated in C minimal medium containing glucose to exponential growth phase (OD_600_, 0.7-0.9) and b-alanine spiked with radiolabeled [1–^14^C]β-alanine (specific activity for both solutes, 50-60 mCi/mmol) together with or without the respective amino acid was added to a final concentration of 30 mM. A sample was taken after 40 minutes by pipetting 500 µl of the culture on a filter. The cells were washed 3 times with 1 ml of C minimal medium and the filter was transferred to a scintillation vial. The filters were completely dried for 30 min at 70°C, before adding 5 ml Ecoscint A liquid scintillation cocktail (National Diagnostics, USA) was added and the radioactivity was measured using a HIDEX 300SL β-particle scintillation detector. For the controls, 500 µl of C Glc with 30 mM β-alanine with and without radiolabeled [1–^14^C]β-alanine were pipetted directly into the scintillation vial and let dried completely at 70°C.

## ACKNOWLEDGEMENTS

This work was supported by grants of the Deutsche Forschungsgemeinschaft (DFG) within the Priority Program SPP1879 (Stu 214-16) (to J.S.) and BBSRC grant BB/Y002644/1 (to R.D.). The funders had no role in study design, data collection, analysis and interpretation, decision to submit the work for publication, or preparation of the manuscript.

## Author contributions

Design of the study: R.W. and J.S. Experimental work: R.W., C.H., and B.H. Data analysis: R.W., C.H., R.D. and J.S. Wrote the paper: R.W., R.D. and J.S.

